# Cancer stem cell plasticity as tumor growth promoter and catalyst of population collapse

**DOI:** 10.1101/018184

**Authors:** Jan Poleszczuk, Heiko Enderling

## Abstract

It is increasingly argued that cancer stem cells are not a cellular phenotype but rather a transient state that cells can acquire, either through intrinsic signaling cascades or in response to environmental cues. While cancer stem cell plasticity is generally associated with increased aggressiveness and treatment resistance, we set out to thoroughly investigate the impact of different rates of plasticity on early and late tumor growth dynamics and the response to therapy. We develop an agent-based model of cancer stem cell driven tumor growth, in which plasticity is defined as a spontaneous transition between stem and non-stem cancer cell states. Simulations of the model show that plasticity can substantially increase tumor growth rate and invasion. At high rates of plasticity, however, the cells get exhausted and the tumor will undergo spontaneous remission in the long term. In a series of *in silico* trials we show that such remission can be facilitated through radiotherapy. The presented study suggests that stem cell plasticity has rather complex, non-intuitive implications on tumor growth and treatment response. Further theoretical, experimental and integrated studies are needed to fully decipher cancer stem cell plasticity and how it can be harnessed for novel therapeutic approaches.

## Introduction

After stem cells have been discovered at the top of the hematopoietic system hierarchy [1], it became apparent that human acute myeloid leukemia is also organized hierarchically. Leukemia is initiated and fueled by a leukemic stem cell that gives rise to transit-amplifying progenitor cells and eventually differentiated cancer cells with limited lifespan [2]. A cellular hierarchy in a tumor and the accompanying stem cell hypothesis has been hailed as a significant breakthrough in the cancer research community, as its concept holds new promises for cancer therapy. If only CSC are uniquely able to initiate, sustain and propagate a tumor, then selective eradication of CSC – however difficult it might be to target them – would be sufficient to cure a cancer [3]. Despite its conceptual beauty, recent reports consolidated the skepticism that stemness might not be a prescribed cell phenotype but a transient state that cells can acquire and discard dependent on the cellular environment and signaling context [4–7]. Then the heterogeneous tumor population becomes a dynamic, moving target that is increasingly difficult to treat [8].

The complex biology of stem and non-stem cancer cells and their interactions with each other as well as with the intra- and extratumoral environment is yet to be fully deciphered experimentally. Inroads have been made to use mathematical and computational models to identify first-order principles and key biological mechanisms in cancer plasticity, from which new actionable hypotheses can be derived [7, 9–12]. Herein we propose an *in silico* agent-based computational model to help decipher parts of the complexity that arises from the myriads of stem and non-stem cancer cell interactions and phenotypic plasticity. Agent-based models are increasingly utilized in theoretical oncology [13–20] to derive emerging population level dynamics from defined single cell properties and their perturbation. Such modeling approach has previously shown that the proliferation capacity (or telomere length [21, 22], or Hayflick limit [23, 24]) of non-stem cancer cell (CC) is a pivotal force in driving tumor evolution [25, 26]. Then, early progenitor cells that adapt a stem cell state confer different kinetic properties including proliferation potential to the new stem cell than de-differentiating cells that are closer to their terminal phenotype.

The role of the cellular hierarchy in a tumor has been widely ignored in the modeling community, such that CC that transition into a CSC have the same properties as the initial population-founding CSC. Thus, in addition to conferring the general CSC traits of (a)symmetric division and longevity, the new CSC gets also bestowed with the historic initial telomere length with the stupendous consequences of increased aggressiveness and treatment failure. Herein we give explicit consideration of the degrading proliferative potential in the cellular hierarchy in a phenotypic plasticity model. We show that phenotypic plasticity promotes early tumor growth; in the long-term, however, plasticity can impede tumor progression and ultimately lead to the collapse of the tumor population. We will use radiotherapy as an example of an external catalyst to eventual complete remission.

## Materials and Methods

### Mathematical model

We adapt an *in silico* agent-based model [26, 27], in which each cancer cell occupies a 10x10 μm grid point on a dynamically expanding 2D lattice [28]. A dynamic computational domain prevents boundary-imposed spatial constrains that may influence outcomes.

The tumor population is divided into cancer stem cells (CSCs) and non-stem cancer cells (CCs) with an individual proliferation capacity, ρ, representative of the telomere length [21, 29]. Telomeres are shortened during mitosis [22, 30] reducing the proliferation capacity in each daughter cell (ρ-1), which is a visualization of the Hayflick limit [24, 31]. CSCs are believed to upregulate telomerase which rebuilds telomeric DNA and thus prevents telomere erosion and confers longevity to the cell [32–35]. We assume a cell with exhausted proliferation capacity (ρ=0) to undergo cell death in the next mitotic attempt as previously assumed [25, 36] without explicit consideration of replicative senescence [18, 37]. In addition to replicative cell death we consider spontaneous cell death in CC with probability α, which is prevented in CSC. Without considering cell plasticity, CC and CSC populations are only connected through asymmetric division of a CSC, when one of the progeny adopts a CC phenotype (Fig. 1A). We denote the probability of symmetric CSC division by p_s_. Plasticity may occur at successful proliferation with probabilities p_d_ (CSC differentiation) and p_dd_(CC de-differentiation). If P_d_=P_dd_=0, plasticity is averted and cells have a persistent phenotype. With p_d_>0 and P_dd_>0, cell phenotypes are plastic and stemness becomes a transient state. We note that stemness is defined by the ability to divide (a)symmetrically and prevention of telomere erosion [34, 38]. Therefore, a CC that adopts a CSC state through dedifferentiation will be equipped with current telomere length, i.e. ρ, which will be bequeathed to subsequent daughter cells (Fig. 1B). This is in stark contrast to previous modeling attempts that in addition to conferring general CSC traits also re-set time and equip new CSC with historic uniform initial telomere length.

**Figure 1.**
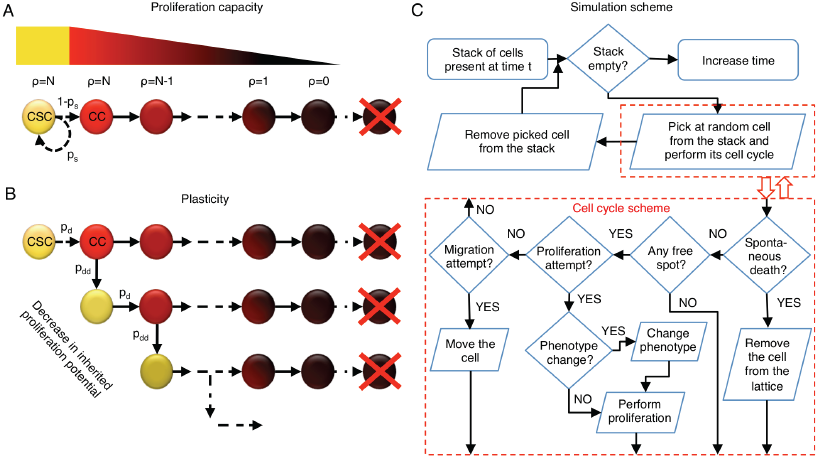
A) Schematic of the proliferation potential erosion in the non-cancer cell (CC) population after asymmetric division of cancer stem cell (CSC) that occurs with probability p_s_. **B)** Schematic of cellular differentiation and de-differentiation (probabilities p_d_ and p_dd_, respectively), in which the proliferation potential is memorized in course of evolution. **C)** Schematic of the simulation procedure and cell cycle evaluations.

At discrete simulation time steps representative of Δt = 1 hour, cell are randomly selected and updated. In case of a CC, spontaneous cell death is considered with probability α. Proliferation and migration of surviving cells are mutually exclusive (p_p_=1/24, i.e. once per day; p_m_=15/24, i.e. 150μm per day) subject to available space in the immediate cell neighborhood. The simulation procedure is summarized in a flowchart in Fig. 1C.

### Tumor morphology analysis

In case of frequent migration events, defining tumor periphery on a two-dimensional lattice is not straightforward, as cells may separate from the main tumor mass. In order to define the tumor periphery we first transform the lattice into binary information (cell present or not). Then we substitute the value of each pixel by the average number of positive pixels in the immediate 8-neighbors Moore neighborhood, and apply an image intensity threshold of 3/8 (this includes vacant sites with more than 2 cancer cells in the neighborhood) and select regions containing more than 10^4^ cells. We define the tumor periphery as the periphery of the selected regions. All transformations are made using MATLAB with Image Processing Toolbox.

Tumor circularity is calculated as

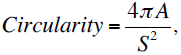

where *A* is the tumor area and S is the perimeter length. For a perfect circle the circularity is equal to 1. Both *A* and *S* are calculated using the *regionprops* function from the MATLAB Image Processing Toolbox.

In order to calculate the CSC fraction in the proximity of the tumor boundary, we dilate the tumor periphery by 20 pixels in four orthogonal directions and select all cells within the dilated area. This procedure selects cells that are within 200 μm in from the tumor boundary.

### Metastatic potential

We estimate the potential for metastatic spread using a virtual 5-well plate, where wells are connected sequentially by a narrow canal of 100 μm width and 400 μm length. Each well is circular with a 50 μm diameter. Each simulation is initiated with a single CSC in the geometric center of the first well. Metastatic spread is simulated until the first CSC enters the last blind-ended canal.

### Radiotherapy

We simulate fractionated radiotherapy of 30 fractions of dose d=2 Gy applied every 24h hours. The dose-dependent surviving fraction SF(d) is calculated using the linear-quadratic formalism

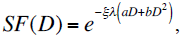

where ξ<1 describes radioresistence of quiescent cell, i.e. for cell with no available space in the neighborhood, and ξ = 1 otherwise, and λ<1 describes increased radioresistance of CSC [39, 40]. For CC, λ = 1. In line with previous estimates we set ξ=0.5, λ = 0.1376, a=0.3859 and b= 0.01148 [40].

## Results and Discussion

We initiate each simulation with a single CSC with proliferation capacity ρ =10 and probability of symmetric division p_s_= 1%, and simulate tumor growth for 720 days. We consider bidirectional plasticity with equal probabilities p_d_ = P_dd_ = 0% (static phenotypes), 0.01%, 0.1%, 1% and 10%. For each set of parameters we performed 100 simulations, and statistical analyses were performed using the Student’s t-test.

### Impact of plasticity on tumor growth characteristics

Phenotypic plasticity increases initial tumor growth rate yielding larger tumors after 720 days than the tumors with static phenotypes (Fig. 2A). However, for larger plasticity (p_d_ = 10%), tumor growth saturates around day 320 keeping the tumor in a dormant state followed by a decrease in total cell number. In contrast to initial growth that is favored by higher plasticity rates, increasing phenotypic plasticity inhibits tumor growth later. This population level behavior mimics the evolution of CSC number (Fig. 2B) and ratio (Fig. 2C). CSC fraction in phenotypic plasticity tumors appears to saturate around the value of 50% for all considered transition rates, but this ratio is only achieved in the considered time frame with p_d_ = P_dd_ =10%.

**Figure 2.**
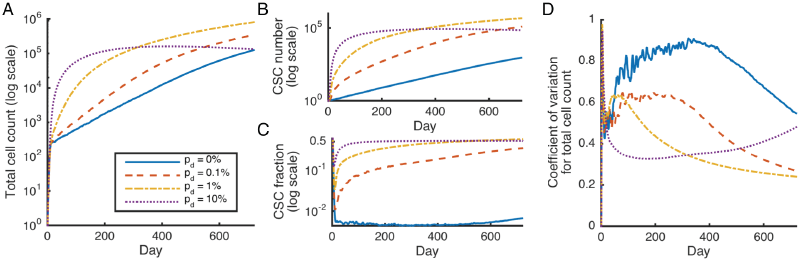
Average total cell count (A), cancer stem cell, CSC, number (B) and fraction (C), and coefficient of variation, i.e. standard deviation/mean, for the total cell count (D) as a function of time for tumor without plasticity (p_d_ = 0%, blue solid curve) and with plasticity probability of 0.1% (red dashed curve), 1% (yellow dot-dashed curve) and 10% (purple dotted curve). Shown are means from 100 simulation runs (only successfully grown tumors are considered). Error bars omitted for clarity.

In addition to increasing the average tumor size during the growth phase, plasticity reduces the amount of variation in tumor size (Fig. 2D). For p_d_ = 1%, standard deviation is only 20% at t=720 days, compared to 3 times larger S.D.=60% for tumors with static phenotypes. The source of variation in tumor size across independent simulation is opportunistic competition for available space between CSC and CC in the tumor interior where most CSC are located without plasticity [19, 25, 36, 41]. Phenotypic plasticity partially averts intratumoral CSC inhibition and prolonged phases of population-level dormancy [42] as new CSC are continuously created at the boundary with fewer spatial constrains.

Interestingly, in case of p_d_= 0.1% one simulate tumor died spontaneously early during development. In that simulation, the initial CSC differentiated before the first stochastic symmetric division event, and all of its CC progenies died before a de-differentiation event. In the Appendix we show analytically that the probability of that event is approximately 0.5% for all considered plasticity rates and, thus, is large enough to manifest in numerical simulations.

To analyze analytically if tumors with phenotypic plasticity can undergo spontaneous remission, we first consider the possible division fates of a CSC with proliferation capacity *i* (CSC_i_). With probability p_d_ the CSC will differentiate before a symmetric division and it will be lost. In case of a de-differentiation event of a CC, the new CSC will have a proliferation capacity ρ < *i* (Fig. 3A). If the CSC divides symmetrically, a new CSC_i_ will be created. In case of asymmetric division we need to follow the fate of the CC progeny. At each iteration, the CC can die spontaneously with probability α. Let us denote by α’ the probability that the newly created CC will die before a proliferation attempt (α’≥α). Then, no new CSC_i_ will be created with probability α’. If the CC divides, a new CSC_i_ is only created in case of a de-differentiation event with probability p_dd_. If after de-differentiation the CSC_i_ undergoes symmetric division, then two new CSC_i_ are be created. This theoretical consideration has four possible outcomes: the number of CSC_i_ 1) decreasees with probability p_1_ = p_d_, 2) remains the same with probability p_2_= (1-p_d_)(1-p_s_)[α’+(1- α’)(1-p_dd_)], 3) increases by one with probability p_3_= (1-p_d_)[p_s_+(1-p_s_)(1-α’)p_dd_(l-p_s_)], and 4) increases by two with probability p_4_ = (1-p_d_)(1-p_s_)(1- α’)p_dd_ p_s_. Hence, the probability P that all CSC_i_ will die after the initial seeding of one CSC_i_ fulfills the relation P = p+qP^2^+(1-p-q)P^3^, where p = p_1_/(p_1_+p_3_+p_4_) and q = p_3_/(p_1_+p_3_+p_4_), from which we obtain that

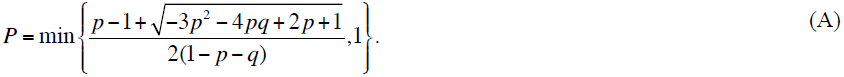

**Figure 3.**
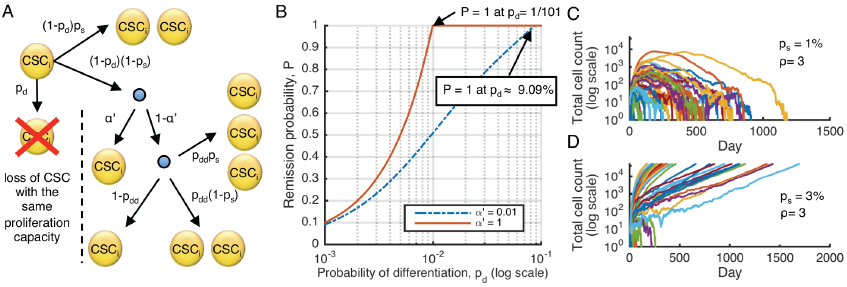
(A) Possible outcomes of the cancer stem cell (CSC) division when one considers survival of CSC with proliferation potential *i* (CSC_i_). (B) Probability of spontaneous tumor remission, P, described by Eqn. (A), for different values of probability that the non-stem cancer cell will die before proliferation attempt, α’, and plasticity event frequencies, p_d_ = p_dd_. (C) Spontaneous remission of all 100 simulated tumors initiated with single CSC equipped with proliferation capacity ρ = 3 and probability of symmetric division p_s_ = 1%. (D) Spontaneous remission of 17 out of 100 simulated tumors initiated with single CSC equipped with proliferation capacity ρ = 3 and probability of symmetric division p_s_ = 3%.

Obviously for p_d_= 0, i.e. no possible differentiation event, we have P = 0, and for α’ = 1 the problem reduces to P = min{(1-q)/q,1}. Importantly, if P = 1, then all CSC will die off regardless of the initial number of CSC. Eqn. (A) contains only one unknown parameter, α’, which depends on the extend of spatial inhibition and the proliferation probability p_p_. However, the probability, p, is an increasing function of α’, and α’ ≥ α. Thus, if a tumor dies spontaneously with probability 1 for α’=α, then it also dies spontaneously with probability one for any larger α’. Similarly, if P < 1 for α’ = 1, then the tumor will not die spontaneously in each single simulation iteration. It is worth to mention that a non-zero value of parameter α is crucial for relating the agent-based model to probability P, as the average time between the CSC proliferation events can grow without bound if α =0, due to intratumoral spatial inhibition.

Fig. 3B shows the probability P of a CSC population vanishing for α’=1 and α’=0.01 if P_d_=P_dd_. All simulated tumors will eventually die off spontaneously for p_d_ values larger than ≈ 9.09%. This explains the previously observed decrease in the average tumor size for p_d_=10% (c.f., Fig. 2A). For p_d_ = 0.1% in the initially presented tumor growth simulations, P~0.1 for α’ = 1, and thus about 90% of tumors will grow successfully.

Let us consider 100 independent simulations initialized with CSC_3_ and p_d_ = P_dd_ = 10%. As predicted by the above theory, all 100 tumors will undergo remission as P = 1 in that case (Fig. 3C). Increase the probability of symmetric division from p_s_=1% to p_s_=3%, only 17 of 100 tumors die out (Fig. 3D). However, the above theory cannot conclusively predict if all of these tumors will eventually die off.

### Impact of plasticity on tumor morphology

Fig.4A shows the largest tumors after simulated 720 days of 100 independent simulations for different plasticity probabilities. Although the average tumor size for p_d_ = 10% is larger than for p_d_=0% (c.f., Fig. 1A), the biggest simulated tumor is smaller (114,950 vs. 283,504 cells). This is a manifestation of the larger coefficient of variation in the static phenotype cohort (c.f., Fig. 1D).

**Figure 4.**
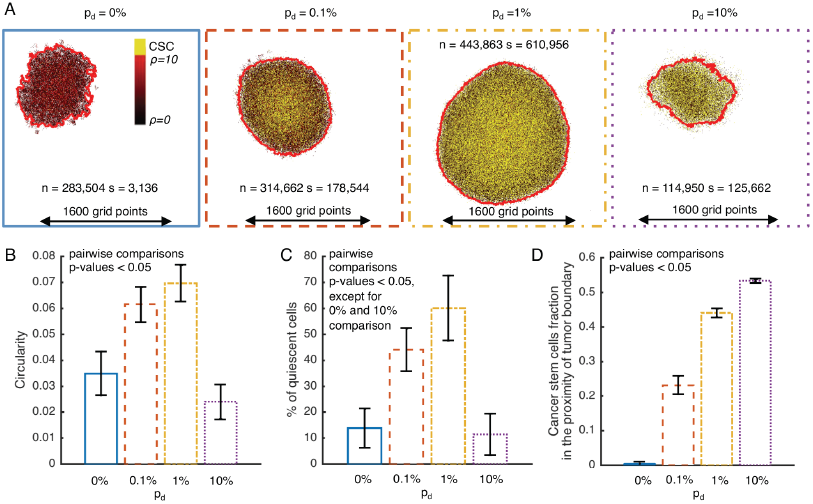
(A) Simulation snapshots of the biggest tumor at t = 720 days for each consider plasticity event probability, s: number of cancer stem cells; n: number of cancer cells. Comparison of circularity (B), percent of quiescent cells (C), and cancer stem cells fraction in 200μm proximity of tumor boundary (D) for different probabilities of plasticity event. Shown are means +− SD from 100 simulation runs (only successfully grown tumors are considered).

The morphology of tumors with static phenotypes is described as self-metastatic [36] with low circularity [43]. Tumors with intermediate plasticity probabilities, p_d_=0.1% and p_d_=1%, feature a regular, almost circular morphology at day 720 as the self-metastatic morphology caused by the spatial inhibition of CSCs is averted by spontaneous CCs de-differentiation at the tumor periphery (Fig. 4B). For larger plasticity probabilities (p_d_ = 10%), clusters of cells at the tumor periphery die off stochastically, which yields a less regular boundary and decreasing tumor circularity.

Low values of plasticity probabilities (pd=0.1% and pd=1%) are associated with a fivefold increase in the fraction of quiescent cells compared to tumors with static phenotypes, arguably at least in part due to the larger size and thus reduced surface-to-volume ratio (Fig. 4C). For p_d_ = 10% the exhaustion and spontaneous death of CC is not effectively filled and thus a larger proportion of cells is actively proliferating, which is comparable to the one in the non-plastic tumor.

A monotonic dependence on plasticity probability is the fraction of cancer stem cells in the proximity of the tumor boundary (Fig. 4D). For p_d_ = 10%, more than 50% of the CSC are close to the tumor boundary, which represents almost the entire cancer stem cell fraction (c.f. Fig. 2C). The prevalence of CSC in the tumor periphery suggests that plasticity may lead to increased potential for metastatic spread.

### Impact of plasticity on invasiveness

For each considered value of plasticity probability we simulated 100 virtual 5-well plate experiments. For p_d_ = 0%, 0.1% and 1%, in all simulations a CSC successfully reached the right boundary of the experimental setup. However, for p_d_= 10% as many as 97 out of 100 simulations ended with spontaneous death of all cancer cells before invading all wells. Time to reach the right boundary was significantly lower in all plastic tumors, with almost 1.5-fold and 2-fold reduction for p_d_ = 0.1% and p_d_ = 1%, respectively, when compared to tumors with static phenotypes (Fig. 5A). However, the invasion speed follows a non-monotonic behavior in plastic populations, as the time needed to reach right boundary for is reduced for p_d_ =1% compared to both p_d_ = 0.1% and p_d_ = 10%, indicating a exhaustion of the population as discussed above.

**Figure 5.**
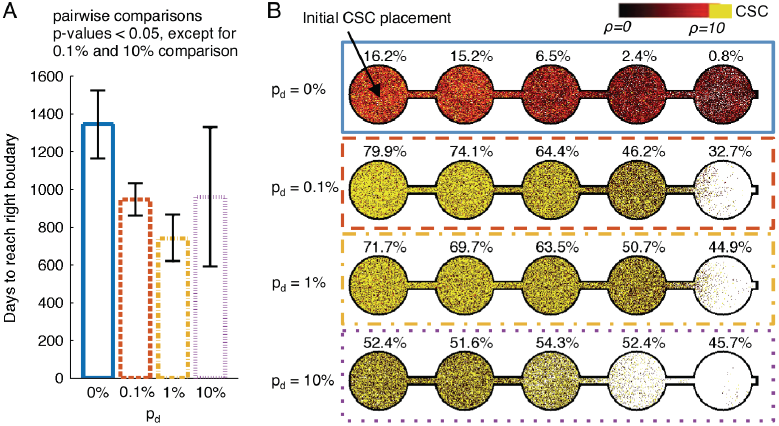
(A) Average ± SD number of days until the right boundary of 5-well plate is reached by cancer stem cell for different probabilities of plasticity event (p_d_). (C) Simulation snapshots at the moment when first cancer stem cell reach the right boundary for different probabilities of plasticity event (p_d_). Percentages above each well indicate the fraction of cancer stem cells within each well.

Visualizations of successful simulations show a plasticity-dependent gradient of cancer stem cell fraction values in subsequent wells (Fig. 5B). With lower values of plasticity the difference between the composition of population in the first and the last well increases, with about 16-fold change for non-plastic tumor. For the largest considered value of p_d_=10%, however, there is no significant difference in composition between the wells.

### Impact of plasticity on the radiation outcome

We simulated the impact of radiotherapy on tumors consisted of 250,000 cells generated for different plasticity probabilities. As expected, the tumor without plasticity responds best to radiotherapy (Fig. 6A), as it has the smallest fraction of CSC with decreased radiosensitivity λ (c.f., Fig. 2C) and small proportion of quiescent cells with decreased radiosensitivity ξ (c.f., Fig. 4C). The tumors with relatively low plasticity, p_d_=0.1% and p_d_=1%, show similar response to radiation, with number of cells reduced about 20 times compared to the pre-treatment state. Tumor with the highest plasticity event probability, p_d_ = 10%, shows an intermediate response, what can be explained by lower fraction of quiescent cells compared to other plastic tumors (Fig. 4C) and larger fraction of stem cells compared to the plasticity free tumor (Fig. 2C). However, in none of simulations tumor was eradicated during the course of treatment.

**Figure 6.**
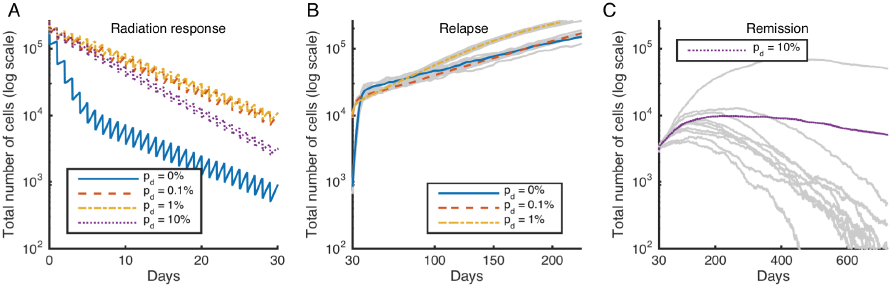
Response to radiotherapy (RT) consisted of 30 x 2Gy dose applied every day of tumors with about 250,000 initial cell count for different probabilities of plasticity event (p_d_). (A) Average radiation response. (B) Regrowth for low p_d_. (C) Remission for p_d_ = 10%. N = 10 simulations.

Simulations of tumor growth following the treatment show that for tumors without and with low plasticity show regrowth to pre-treatment cell counts within 200 days after the onset of treatment (Fig. 6B). However, all of the simulated tumors with p_d_ = 10% undergo spontaneous remission (Fig. 6C) within two years after transient regrowth. Radiotherapy on highly plastic tumors may serve as an accelerator of CSC exhaustion due to frequent differentiation events with subsequent loss of proliferation potential.

## Conclusions

The observation of cancer stem cell plasticity has lead to the general belief of more aggressive tumor growth and reduced treatment response [12]. We introduced an *in silico* agent-based model of cancer stem cell driven tumor growth to study the impact of different rates of phenotypic plasticity on tumor growth, morphology, invasion and treatment response. Simulations of our model show that plasticity accelerates early tumor growth, as prolonged phases of tumor dormancy due to intratumoral competition [42] is averted by cells on the periphery acquiring stem cell traits and propagating the tumor population. Whilst tumors with fixed phenotypes show a gradual increase in CSC over time, phenotypic plasticity yields relatively constant population fractions, with CSC predominantly located at the tumor boundary – thereby facilitating invasion and metastatic spread. The explicit consideration of the cellular hierarchy within a tumor and the accompanying reduction of proliferation potential, however, offer a previously unappreciated aspect of CSC plasticity. Transitions between stem and non-stem cancer cell states at high frequencies yield a reduction of proliferation potential in each cell, thereby reducing the lifespan of each daughter cell and inevitably cell death. This may lead to population level dormancy and ultimately collapse of the tumor with complete remission.

Mathematical and computational models, by virtue of their very purpose, are subject to gross oversimplifications of reality [44]. However, they may provide interesting and non-intuitive insights into the tumor progression dynamics, including novel discussion points in understanding cancer stem cell plasticity. Further theoretical, experimental and integrated studies are needed to fully decipher cancer stem cell plasticity and how it can be harnessed for novel therapeutic approaches.

## Acknowledgments

JP would like to thank the Foundation for Polish Science for partial support.

## Appendix Probability of early tumor death event

We set on to approximate analytically the probability that the initial CSC with proliferation capacity ρ will differentiate before first symmetric division and all of its non-stem progenies will die before any de-differentiation event, which we denote as D_ρ_. We consider a very early stage of tumor development and, thus, we can neglect space inhibition (we have large migration probability of 15/24). The probability that non-stem cell with given proliferation potential, ρ, and all its progenies won’t de-differentiate before dying off, ω_ρ_, follows recurrence relation

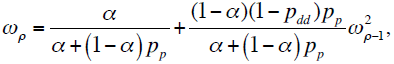

with ω_0_ = 1 – p_p_(1-α)p_dd_/(α+p_p_(1-α)). The probability that the CSC will undergo differentiation before first symmetric division is equal to p_d_/((1-p_d_)p_s_+p_d_) and the number of non-symmetric divisions follows geometric distribution with probability of success p = (1-p_d_)p_s_+p_d_. Thus, we obtain the following expression for D_ρ_

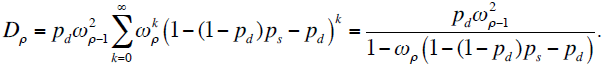

In Fig. A1 we plotted D_ρ_ under the assumption that p_d_=p_dd_ and for parameters considered in the main text. We see that for all considered plasticity event probabilities the probability of early tumor death is close to 0.5%. It is worth to notice that if spatial inhibition will increase value D_ρ_ more for lower probabilities of plasticity event, as on average more non-stem progenies are created before differentiation event.

**Figure A1.**
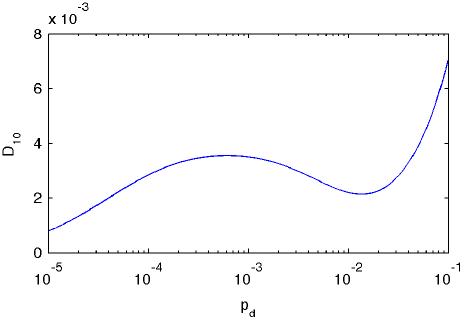
Probability of early tumor death event for different plasticity event probability and for parameters considered in the main text.

